# Heightened virulence of *Yersinia* is associated with decreased function of the YopJ protein

**DOI:** 10.1101/2021.03.22.436461

**Authors:** Chris A. Mares, Fernando P. Lugo, Mohammad Albataineh, Beth A. Goins, Irene G. Newton, Ralph R. Isberg, Molly A. Bergman

**Author notes:** Department of Life Sciences, Texas A&M University-San Antonio, San Antonio, Texas, United States of America (Chris A. Mares).

## Abstract

Despite the maintenance of YopP/J alleles throughout the human-pathogenic *Yersinia* lineage, the benefit of YopP/J-induced phagocyte death for *Yersinia* pathogenesis in animals is not obvious. To determine how sequence divergence of YopP/J has impacted *Yersinia* virulence, we examined protein polymorphisms in this Type III secreted effector protein across 17 *Yersinia* species, and tested the consequences of polymorphism in a murine model of sub-acute systemic yersiniosis. Our evolutionary analysis revealed that codon 177 has been subjected to positive selection - the *Y. enterocolitica* residue had been altered from a leucine to a phenylalanine in nearly all *Y. pseudotuberculosis* and *Y. pestis* strains examined. Despite being a minor change, as both leucine and phenylalanine have hydrophobic side chains, reversion of YopJ^F177^ to the ancestral YopJ^L177^ variant yielded a *Y. pseudotuberculosis* strain with enhanced cytotoxicity towards macrophages, consistent with previous findings. Surprisingly, expression of YopJ^F177L^ in the mildly attenuated *ksgA^-^* background rendered the strain completely avirulent in mice. Consistent with this hypothesis that YopJ activity indirectly relates to *Yersinia* pathogenesis *in vivo*, *ksgA^-^* strains lacking functional YopJ failed to kill macrophages but actually regained virulence in animals. Also, treatment with the anti-apoptosis drug suramin prevented YopJ-mediated macrophage cytotoxicity and enhanced *Y. pseudotuberculosis* virulence *in vivo*. Our results demonstrate that *Yersinia*-induced cell death is detrimental for bacterial pathogenesis in this animal model of illness, and indicate that positive selection has driven YopJ/P and *Yersinia* evolution towards diminished cytotoxicity and increased virulence, respectively.

## Introduction

Bacterial pathogens induce host cell death by various mechanisms, and with downstream consequences that seem counterintuitive (1). At face value, death of host cells, particularly cells of the innate immune system, should cripple the host immune response and allow pathogens to escape clearance. However, the opposite often occurs – induction of host cell death can induce a host response that leads to pathogen removal (1). Mechanisms underlying this outcome are incompletely understood, but likely include removing niches for intracellular replication, or enhancing recruitment of highly activated phagocytes that then ingest and degrade the offending bacteria. Both host and bacterial factors are necessary for pathogen-induced host cell death – caspase-1 being a notable player during macrophage pyroptosis (2) and required for host protection against pyroptosis-inducing pathogens (3, 4). Thus, pathogen-triggered cytotoxicity is a host defense strategy. But why do pathogens maintain the genes encoding the pro-cytotoxicity factors in their genomes? The existence of such genes throughout the bacterial kingdom suggests that host cell death plays an important role in persistence of individual species.

Three species of mammalian-pathogenic *Yersinia,* the gastroenteritis-causing *Y. enterocolitica* and *Y. pseudotuberculosis* and the plague-causing *Y. pestis*, have maintained such a pro-cytotoxicity factor throughout their evolutionary history, in spite of significant genomic decay (5, 6). YopJ (so-called in *Y. pestis* and *Y. pseudotuberculosis*, called YopP in *Y. enterocolitica*) is one of the 6-8 Type 3 secretion system (T3SS) effector proteins (*Yersinia* outer proteins, Yops) injected by the bacteria into host cells (7-9). YopJ has been described as having ubiquitin-like protein protease (10), deubiquitinase (11, 12) and acyl transferase activity (13, 14), with the latter requiring the host cell co-factor inositol hexaskisphosphate (15). In macrophages, YopJ activity inhibits MAPK and NFκB signaling pathways, which in conjunction with TLR4 signaling results in host cell death (reviewed in (16)), and recent evidence demonstrates that YopJ-induced phagocyte death occurs by a mechanism distinct from both apoptosis and pyroptosis but linked to necrosis(17). Regardless of mechanism, the outcome is clear – YopJ has profoundly negative effects on macrophage viability.

However, exactly what YopJ does for *Yersinia in vivo* is unclear. YopJ seems minimally important for virulence, at least compared to most of the other Yops, as YopJ^-^ *Y. pestis,* YopJ^-^ *Y. pseudotuberculosis*, and YopP^-^ *Y. enterocolitica* all display near-wildtype levels of virulence in various mouse models of acute illness (18-23). In contrast, there is evidence indicating that YopJ activity may actually diminish *Yersinia* virulence, as different isoforms of YopP/J can alter the degree of macrophage death and mouse illness following exposure to *Yersinia*. *Y. pseudotuberculosis* or *Y. pestis* expressing a hypersecreted variant of YopP showed enhanced cytotoxicity towards cultured macrophages but diminished virulence in mice (24, 25). Moreover, *Y. pestis* strain KIM also encodes a hypercytotoxic YopJ variant, which when expressed in *Y. pseudotuberculosis* induced more macrophage cell death due to enhanced IKKβ binding and thus reduced NFκB signaling; the impact of this variant upon *Y. pseudotuberculosis-*caused illness in mice was not reported (26).

It is unknown if changes in YopP/J sequences occurred as a consequence of Darwinian evolution, but given that host-pathogen interactions can exert strong natural selection pressure upon both organisms (27), it seems likely that the YopJ-target protein interactions have been guided by evolutionary pressure. Evidence of selection at the codon level can be detected via phylogenetic analyses across multiple lineages to detect patterns of protein polymorphisms relative to divergence (28, 29). The advantage of this approach is that it can reveal loci affected by positive selection, even if the selection pressure is obscure. Such analyses of bacterial pathogen effector proteins have revealed signatures of molecular evolution, with individual codons showing evidence of directional selection (30, 31).

To address the hypothesis that YopP/J has been subjected to directional selection, we analyzed 17 sequences from multiple strains of each human-pathogenic *Yersinia* species for evidence of microevolution, and evaluated how the different isoforms impacted the outcome of systemic sub-acute yersiniosis in a small animal model. Our studies reveal that some YopJ residues differing between the species have been subjected to evolutionary pressure. Reversion of a positively selected residue to the ancestral one yielded a YopJ isoform with enhanced cytotoxicity, but one that markedly attenuated *Y. pseudotuberculosis* virulence in animals. Conversely, loss of YopJ due to catalytic inactivation or deletion rendered *Y. pseudotuberculosis* strains non-cytotoxic but hypervirulent. Chemical inhibition of macrophage death during *in vivo* infection also enhanced *Y. pseudotuberculosis* virulence. Our results indicate that YopJ has evolved towards diminished ability to induce macrophage cell death and that YopJ-induced death is detrimental for *Y. pseudotuberculosis* infection. This may also suggest that some level of YopP/J-induced death is beneficial to the genus, perhaps by allowing persistence in an unknown reservoir.

## Results

### YopJ has been positively-selected in *Y. pestis* and *Y. pseudotuberculosis* for reduced cytotoxicity and enhanced virulence

The *yopP/J* gene has been maintained in the genomes of all strains of human pathogenic *Yersinia* species sequenced to date (5, 32, 33). This conservation is suggestive of an important role for the YopJ protein in the natural setting. Although YopJ homologs are found in the genomes of all 3 human pathogenic *Yersinia* species, sequence conservation differs across the length of the molecule. Some of the sequence polymorphisms group *Y. pestis* and *Y. pseudotuberculosis* to the exclusion of *Y. enterocolitica* (the expectation), while some other polymorphisms do not cleanly follow the predicted phylogeny of the organisms (**Figure 1A**). We investigated whether or not these divergent residues show evidence of positive selection using a likelihood ratio test comparing support in the data for various models of evolution (either allowing for selection or imposing a nearly neutral model of evolution) (34, 35). This analysis identified sites under positive selection (that is, ancestral sequence reconstruction and the number and kind of differences at each residue suggested that dN/dS >1, **Supplementary Tables 1,2**) and included residue 177, which is a leucine in *Y. enterocolitica* and *Y. pestis* KIM but a phenylalanine in the other *Y. pestis* homologs examined and all the *Y. pseudotuberculosis* YopJ homologs (**Figure 1B**). Interestingly, the YopJ^F177L^ isoform was recently described to have enhanced activity relative to the YopJ^F177^ protein, promoting phosphorylation of IκB-α, increased binding to IKKβ, and causing enhanced macrophage cytotoxicity (26).

**Figure 1.**
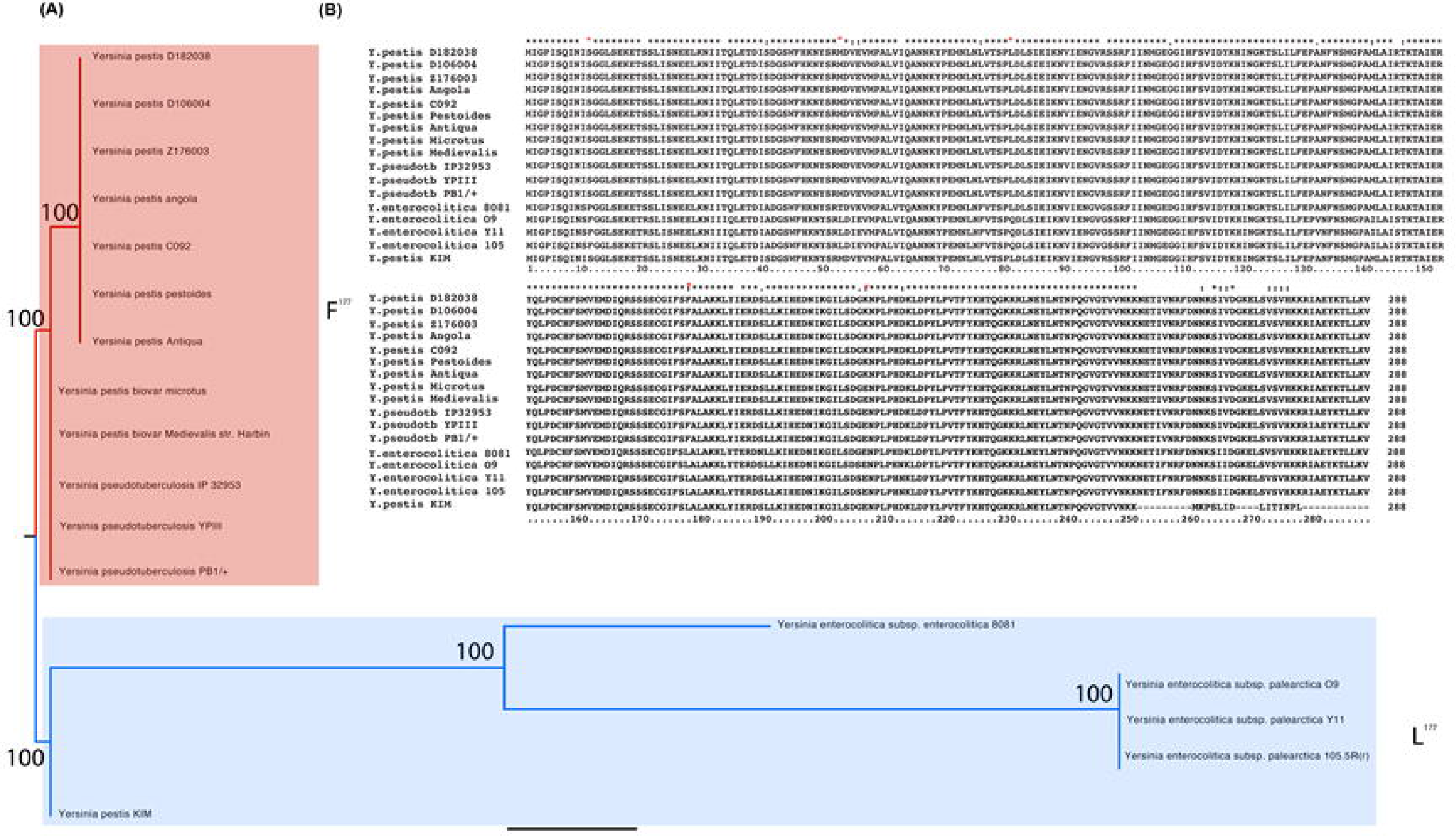
YopP/J is divergent between *Yersinia* species and shows evidence of positive selection. (A) The phylogeny of three *Yersinia* species (*Y. enterocolitica,* 3 strains, *Y. pseudotubercolosis,* 2 strains, and *Y. pestis,* 9 strains) generated using the YopJ primary sequences. Bootstrap support based on 1000 replicates. Codon 177 was identified as under positive selection in a Naive empirical Bayes (NEB) analysis and amino acid polymorphism at that site is mapped onto the phylogeny of these *Yersinia* strains (“L” in blue and “F” in red). (B) Alignment of the relevant region of YopP/J homologs from these 14 *Yersinia* strains from the three human pathogenic species. YopP/J was identified as having experienced positive selection at residue 177.

The existence of a hypercytotoxic YopJ variant allowed us to query if enhanced YopJ activity altered *Y. pseudotuberculosis* virulence in a small animal model of systemic subacute illness. Naïve mice rapidly succumb to infection with virulent Y. pseudotuberculosis. Therefore, in order to better understand the phenotypes in this study we utilized a strain deficient in KsgA that had previously been shown to have a slower replication rate than wild-type Y. pseudotuberculosis due to a loss of demethylation of 16s rRNA (Mecsas, Bilis, and Falkow, 2001, Mangat and Brown). We, and others, have also documented that this strain is attenuated in vivo (Mecsas Bilis, and Falkow 2001, Bergman, et al. 2009). Furthermore, we utilized the intravenous route of infection in order to investigate the role of YopJ after *Y. pseudotuberculosis* has disseminated to distal target tissues. As reported previously, YopJ^F177L^ shows increased cytotoxicity for macrophages relative to the YopJ^F177^ isoform (Fig. 2A). Using the attenuated strain as our baseline background strain (*ksgA^-^*), we observed that *ksgA^-^* bacteria expressing YopJ^F177L^ were more attenuated for virulence than the parental *ksgA^-^* strain, as demonstrated by the 100% survival rate of mice exposed to the *ksgA*^-^ *yopJ^F177L^* strain (**Figure 2B**). Increasing the inoculum dose did not reverse the strain’s complete attenuation (data not shown). These results indicate that the YopJ^F177L^-containing isoform, attenuates virulence in the derived *Y. pseudotuberculosis* strains, correlating with the enhanced cytotoxicity conferred by this variant.

**Figure 2.**
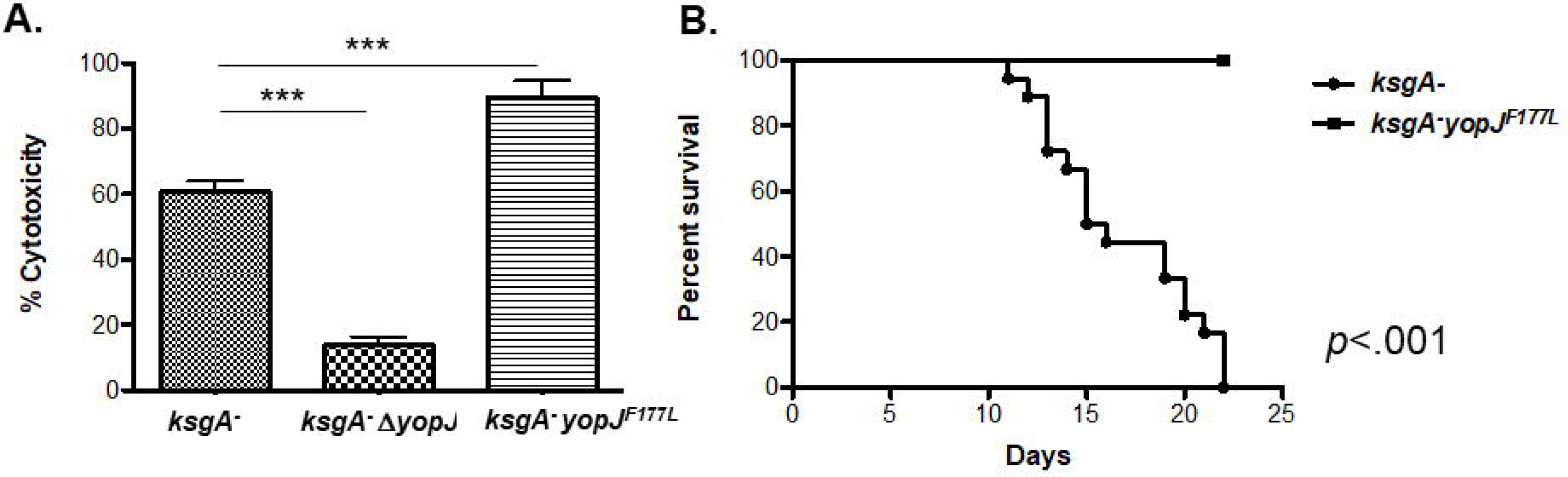
Ancestral version of YopP/J confers enhanced macrophage death and attenuated virulence. (A) Bone marrow derived macrophages were exposed to either *ksgA^-^*, *ksgA^-^*⊗*yopJ*, or *ksgA^-^ yopJ^F177L^ Y. pseudotuberculosis* at an MOI of 100:1. LDH release was measured to determine the cytotoxicity of these strains in macrophages. (B) C57BL/6 mice were intravenously challenged with 1x10^3^ CFU of either *ksgA^-^* (*n*=12) or *ksgA^-^ yopJ^F177L^* (*n*=11) *Y. pseudotuberculosis*. Morbidity and mortality were followed after challenge and data shown represents the percent survival of each group. *In vivo* data is compiled from two independent experiments. Student’s t test was used to analyze BMDM infections. The Kaplein-Meier method was used to generate survival curves and the log-rank test was used to calculate the significance (\**p*<.05,\*\**p*<.01, \*\*\**p*<.005).

### *Y. pseudotuberculosis* strains lacking functional YopJ display increased virulence in vivo

Given that a hypercytotoxic YopJ isoform attenuated *Y. pseudotuberculosis* illness in mice, we predicted that the absence of YopJ would enhance the organism’s virulence. To test this hypothesis, we constructed an isogenic *yopJ* deletion strain in the *ksgA^-^* background, and confirmed that *ksgA^-^* Δ*yopJ* bacteria were non-cytotoxic for bone-marrow-derived macrophages (**Figure 3A**). Consistent with the negative correlation between cytotoxicity and virulence, mice inoculated with the *ksgA^-^* Δ*yopJ* strain died significantly sooner than those challenged with the *ksgA^-^* strain (**Figure 3B**), with kinetics similar to the parental *ksgA^+^* strain (36). Confirming that the enhanced virulence phenotype of the *ksg^-^* Δ*yopJ* strain was due to the *yopJ* deletion, a strain rescued for the genomic Δ*yopJ* lesion by allelic exchange (*yopJ^repaired^*) caused a mortality rate identical to the parental *ksgA^-^* strain in animals (**Supplementary Figure 1**).

**Figure 3.**
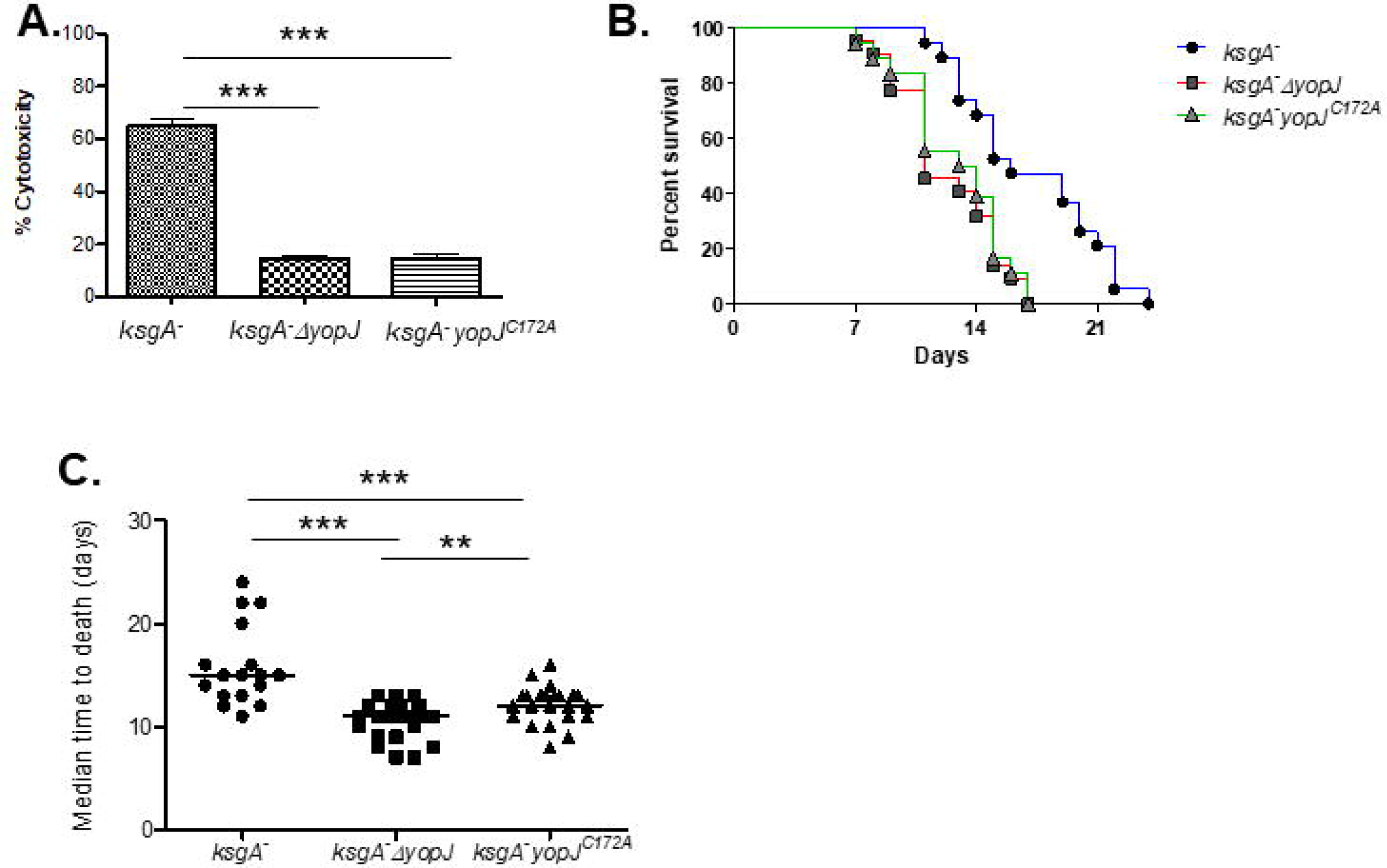
Loss of YopJ reverses the attenuation phenotype of *ksgA^-^ Y. pseudotuberculosis*. (A) Bone marrow derived macrophages were infected with either *ksgA^-^*, *ksgA^-^* Δ*yopJ*, or *ksgA^-^ yopJ^C172A^ Y. pseudotuberculosis* at an MOI of 100:1 as described in the Methods section. One of four independent experiments is shown. LDH release was measured to determine the cytotoxicity of these strains in macrophages. (B) C57BL/6 mice were intravenously challenged with 1x10^3^ CFU of either *ksgA^-^* (*n*=19), *ksgA^-^* Δ*yopJ* (*n*=22), or *ksgA^-^yopJ^C172A^* (*n*=18) *Y. pseudotuberculosis* and monitored for survival (*ksgA^-^* vs. *ksgA^-^ΔyopJ*, p<.001; *ksgA^-^* vs. *ksgA^-^yopJ^C172A^*, p<.001, *ksgA^-^ΔyopJ* vs*. ksgA^-^yopJ^C172A^*, not sig.). (C) The median time until death was also assessed for each experiment. *In vivo* data is compiled from 4 independent experiments. The Student’s t test was used to analyze BMDM infections. The Kaplein-Meier method was used to generate survival curves and the log-rank test was used to calculate the significance. The Mann-Whitney U test was used to compared MTD data (\**p*<.05,\*\**p*<.01, \*\*\**p*<.005).

The enzymatic activity of *Y. pseudotuberculosis* YopJ has been mapped to a triad of residues – histidine 109, glutamate 128, cysteine 172 (10). We asked if the virulence phenotype of a strain carrying the catalytic *yopJ^C172A^* mutation mimicked that observed with the isogenic *ΔyopJ* strain. Similar to previous reports, the *ksgA^-^yopJ^C172A^* strain was unable to kill cultured macrophages (**Figure 3A**) (37). Like *ksgA^-^ΔyopJ* bacteria, *ksgA^-^ yopJ^C172A^ Y. pseudotuberculosis* displayed increased virulence in animals relative to the parental strain (median time to death day 11 versus 12, respectively). Interestingly, *ksgA^-^yopJ^C172A^* bacteria and *ksgA^-^*Δ*yopJ* bacteria consistently displayed similar virulence and median times to death in multiple mouse survival assays (**Figure 3B and 3C**). Taken together, these results indicate that either the absence of YopJ or the presence of inactive YopJ renders an attenuated strain of *Y. pseudotuberculosis* more virulent *in vivo*.

### The YopJ^F177L^ isoform attenuates bacterial burden and increases host cell death *in vivo*

To further understand the pathogenesis of the different YopJ-expressing strains *in vivo,* we examined the kinetics of bacterial colonization in the spleen and liver post-challenge, examining burden at days 1, 3, 9, and 11 post-challenge. This analysis revealed two trends. First, at later time points, mice exposed to the *ksgA^-^ yopJ^F177L^* strains carried lower burdens of splenic and hepatic bacteria than mice exposed to the parental *ksgA^-^, ksgA^-^*Δ*yopJ,* and *ksgA^-^yopJ^C172A^* strains (**Figure 4A, B**). Second, the number of YopJ^F177^ and YopJ-deficient bacteria (both YopJ^-^ and YopJ^C172A^ mutants) generally increased over the 11 day time course in target organs (**Figure 4A, B**), whereas the *ksgA^-^yopJ^F177L^* bacteria were either slower to accumulate in tissues (liver, **Figure 4A**) or diminished during this time frame (spleen, **Figure 4B**). These results are consistent with YopJ triggering host immune responses that inhibit bacterial replication and/or remove bacteria from the tissue, while the absence of YopJ activity allows the bacteria to replicate unchecked in tissues.

**Figure 4.**
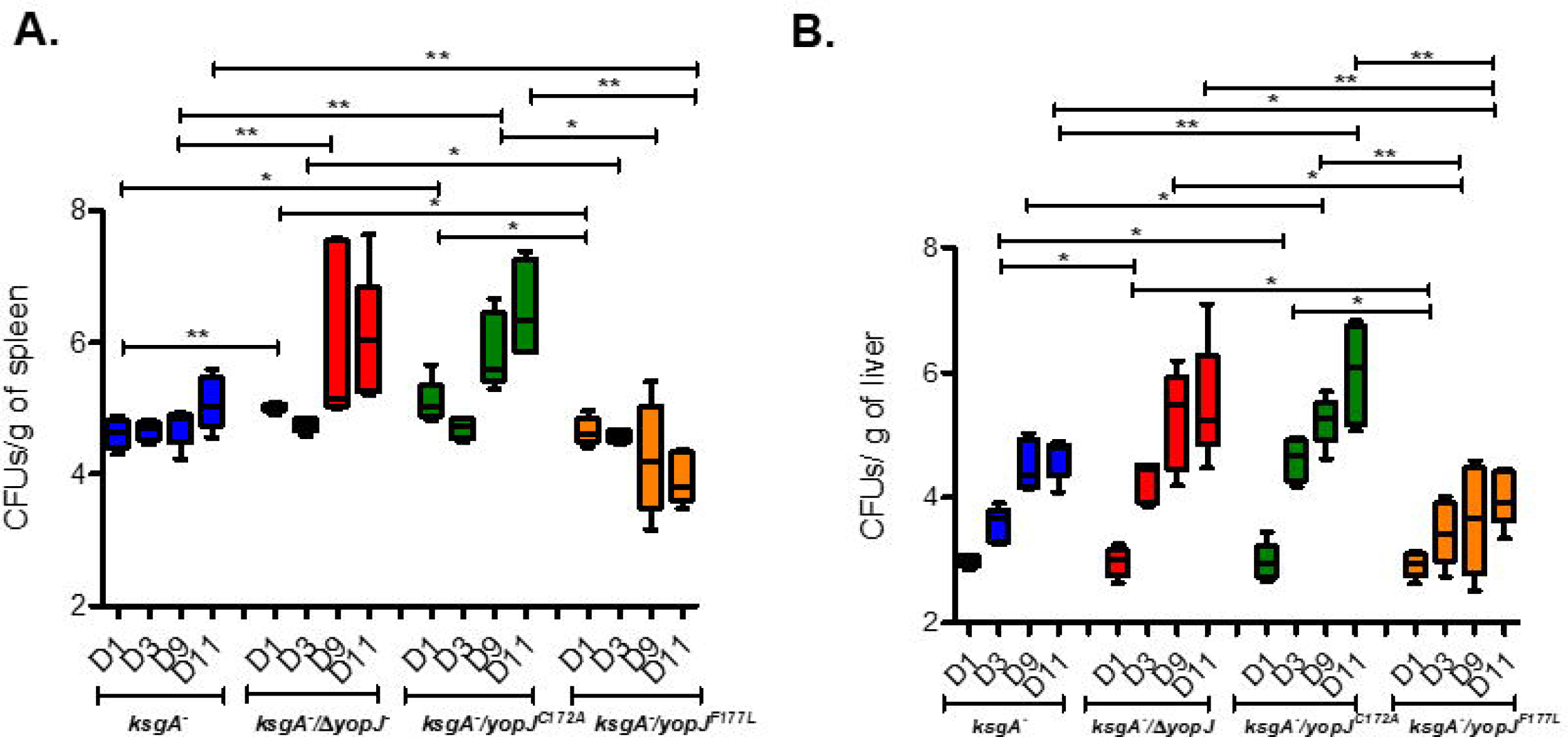
*Y. pseudotuberculosis* expressing YopJ^F177L^ show attenuated colonization *in vivo*. (A) Liver and (B) spleen were isolated at multiple time points from C57BL/6 mice after post-challenge (1.66∼1.94x10^3^ CFU) with *ksgA^-^*, *ksgA^-^*Δ*yopJ*, *ksgA^-^ yopJ^C172A^*, or *ksgA^-^yopJ^F177L^ Y. pseudotuberculosis*. The tissues were homogenized, serially diluted and plated on LB plates. Burdens recovered in each tissue from each mouse were above the limit of detection. Data shown is from 5 mice/group and is representative of two independent experiments. The Mann-Whitney U test was used to compare CFU data (\**p*<.05,\*\**p*<.01, \*\*\**p*<.005).

In addition to causing macrophage cytotoxicity, YopJ can also inhibit macrophage production of cytokines (38-41), such that the increased mortality of mice exposed to *ksgA^-^* bacteria carrying either a null or inactive *yopJ* allele could result from a massive overproduction of cytokines, also known as a cytokine storm (42-44). However, we did not find any evidence indicating that the absence of functional YopJ triggers a mortality-inducing cytokine storm *in vivo* (**Supplementary Figure 2**).

To determine if cell death levels *in vivo* recapitulated what we had observed with cultured macrophages exposed to the different *Y. pseudotuberculosis* strains, we used the TUNEL assay to detect dead mammalian cells in tissue sections from colonized mice. Mice were sacrificed at day 3 post-inoculation for cell death evaluation to ensure equivalent burden levels between mice inoculated with the different strains (as observed in **Figure 4**). This analysis revealed that at 3 days post infection, spleens containing *ksgA^-^* (WT YopJF177) or *ksgA^-^yopJ^F177L^* bacteria had increased levels of TUNEL^+^ cells as compared to spleens containing the *ksgA^-^*ΔyopJ bacteria (**Figure 5** **A-D)**, although the strain expressing catalytically dead YopJ did not perfectly phenocopy the *ksgA^-^* Δ*yopJ* strain (**Figure 5D**). These results indicated that the increased survival of mice infected with *ksgA^-^yopJ^F177L^* was significantly correlated with the amount of TUNEL staining present in their spleens at day 3 post infection. Conversely, a decrease in the amount of TUNEL staining present in the spleens at this time point was associated with the early death of mice infected with either *ksgA^-^*Δ*yopJ* or with *ksgA^-^ yopJ^C172A^*.

**Figure 5.**
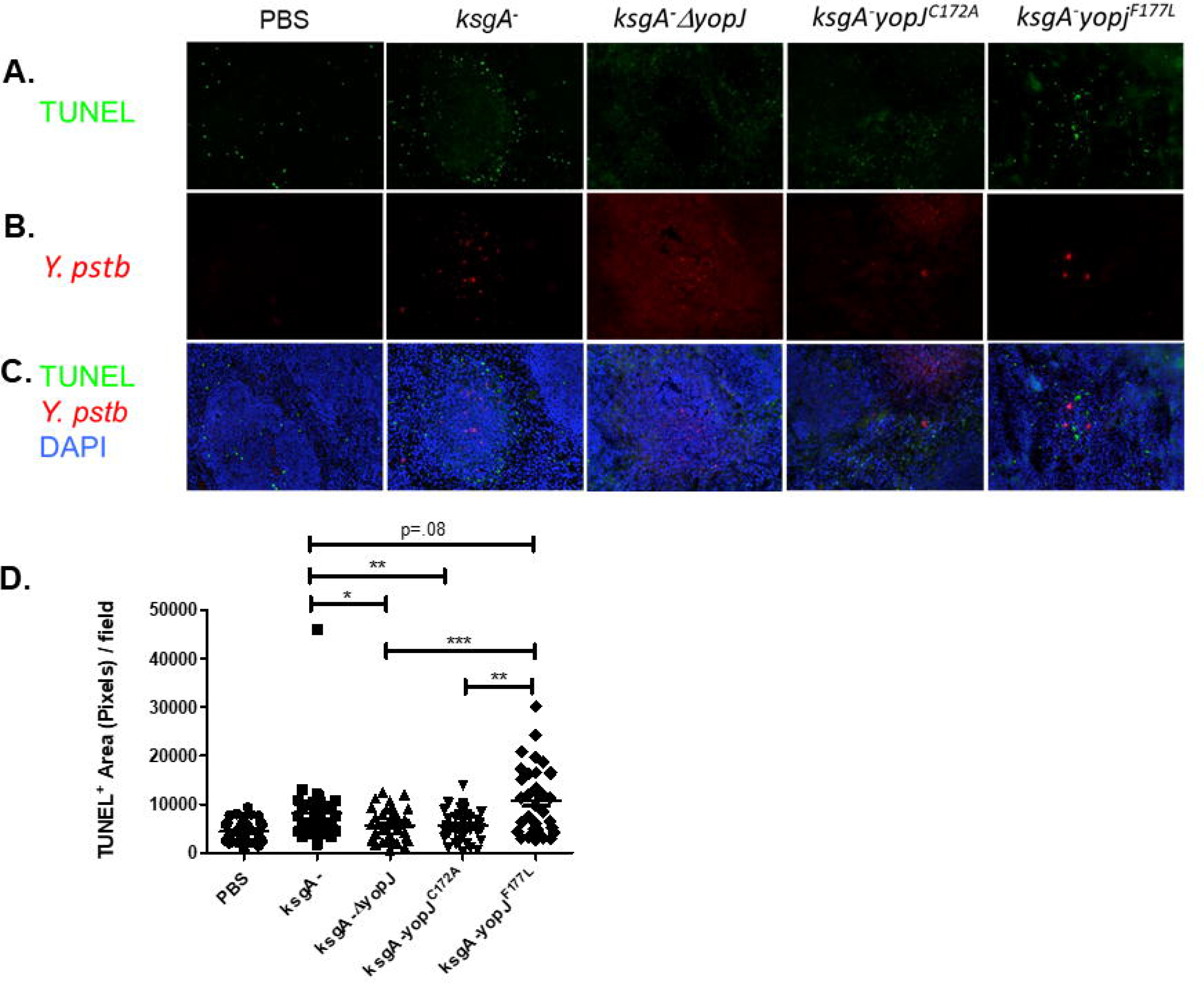
Ancestral YopP/J variant causes increased levels of cytotoxicity *in vivo*. 6-8 week old female C57BL/6 mice were infected intravenously (1∼1.26x10^3^ CFU) with strains of *Y. pseudotuberculosis*expressing *ksgA^-^*, *ksgA^-^*Δ*yopJ*, *ksgA^-^ yopJ^C172A^*, or *ksgA^-^ yopJ^F177L^*. **A-C.** Spleens were removed at day 3 post infection snap frozen in OCT for TUNEL staining. Row A is showing TUNEL only images at 200x and row B is showing staining of representative images for *Y. pseudotuberculosis* (Rhodamine Red X). Row C is depicting the merged images with DAPI (blue), TUNEL (FITC) and *Y. pseudotuberculosis* (Rhodamine Red X) at a total magnification of 200x. **D.** Random images were captured (6 from each spleen, 3 mice per group per experiment) and TUNEL+ area (total pixels) per field was quantified using ImageJ software. 5D shows pooled results obtained from two independent experiments. The Mann-Whitney U test was used to compare TUNEL data (\**p*<.05,\*\**p*<.01, \*\*\**p*<.005).

### Suramin inhibits Yersina-induced macrophage cytotoxicity and enhances virulence *in vivo*

Our results suggest that YopJ-induced cytotoxicity is detrimental to *Y. pseudotuberculosis* pathogenesis, such that enhancing or inhibiting of host cell death *in vivo* should impair or promote *Y. pseudotuberculosis* virulence, respectively. To test this hypothesis, we first considered if artificially increasing macrophage death in mice exposed to YopJ-null *Y. pseudotuberculosis* would rescue the mice from illness. Clodronate-containing liposomes are commonly used to deplete macrophages *in vivo –* depletion results because the clodronate induces global macrophage death (45); we confirmed that mice exposed to clodronate liposomes showed massive levels of apoptotic cells in tissues (data not shown). We inoculated mice with *ksgA^-^*Δ*yopJ Y. pseudotuberculosis* and 6 hours later, delivered a one-time injection of clodronate liposomones. The liposome-treated animals showed a slightly increased rate of mortality as compared to mock-treated animals (**Supplementary Figure 3**), suggesting that the global depletion of macrophages masked any consequence of increased macrophage apoptosis.

We next attempted to diminish host cell death levels *in vivo*, to determine if this would enhance *Y. pseudotuberdulosis* virulence. We turned to the apoptosis inhibitor, suramin, which is a polysulfonated urea derivative capable of inhibiting death receptor-induced apoptosis and apoptosis-mediated liver damage (46, 47). Suramin blocked Yersina induced death of cultured macrophages in a dose-dependent manner (**Figure 6A**), at the same levels previously shown to block hepatic cell cytotoxicity induced by death receptors CD95, TRAIL-R1 and TRAIL-R2 (46). To evaluate how suramin treatment affected *Y. pseudotuberculosis* virulence *in vivo*, mice were inoculated with bacteria, then injected with suramin 30 minutes to 1 hour post-inoculation. Suramin had no effect on naïve mice, but dramatically increased mortality of mice exposed to *ksgA^-^* bacteria (**Figure 6B**). This effect was YopJ-independent, however, as mice inoculated with *ksgA^-^* Δ*yopJ* bacteria and subsequently injected with suramin showed a similar increase in time-to-death (**Figure 6B**). Strikingly, gross pathological differences were apparent after examination of H&E stained liver sections taken from *ksgA^-^*-infected mice treated either with suramin or PBS. The suramin-treated/infected mice had significantly fewer microabcesses visible in liver sections when compared to PBS-treated/infected mice (**Figure 6C-E**). Confirming that suramin successfully impaired host cell death, liver sections taken from suramin-treated mice at 3 days post-bacterial inoculation showed significantly fewer TUNEL+ cells within abcesses than untreated mice (**Figure 6F-H**). These results demonstrate that inhibition of host cell death dramatically worsens the survival outcome for mice experiencing *Y. pseudotuberculosis-*caused illness, and strongly supports the premise that host cell death protects the host and checks the pathogen.

**Figure 6.**
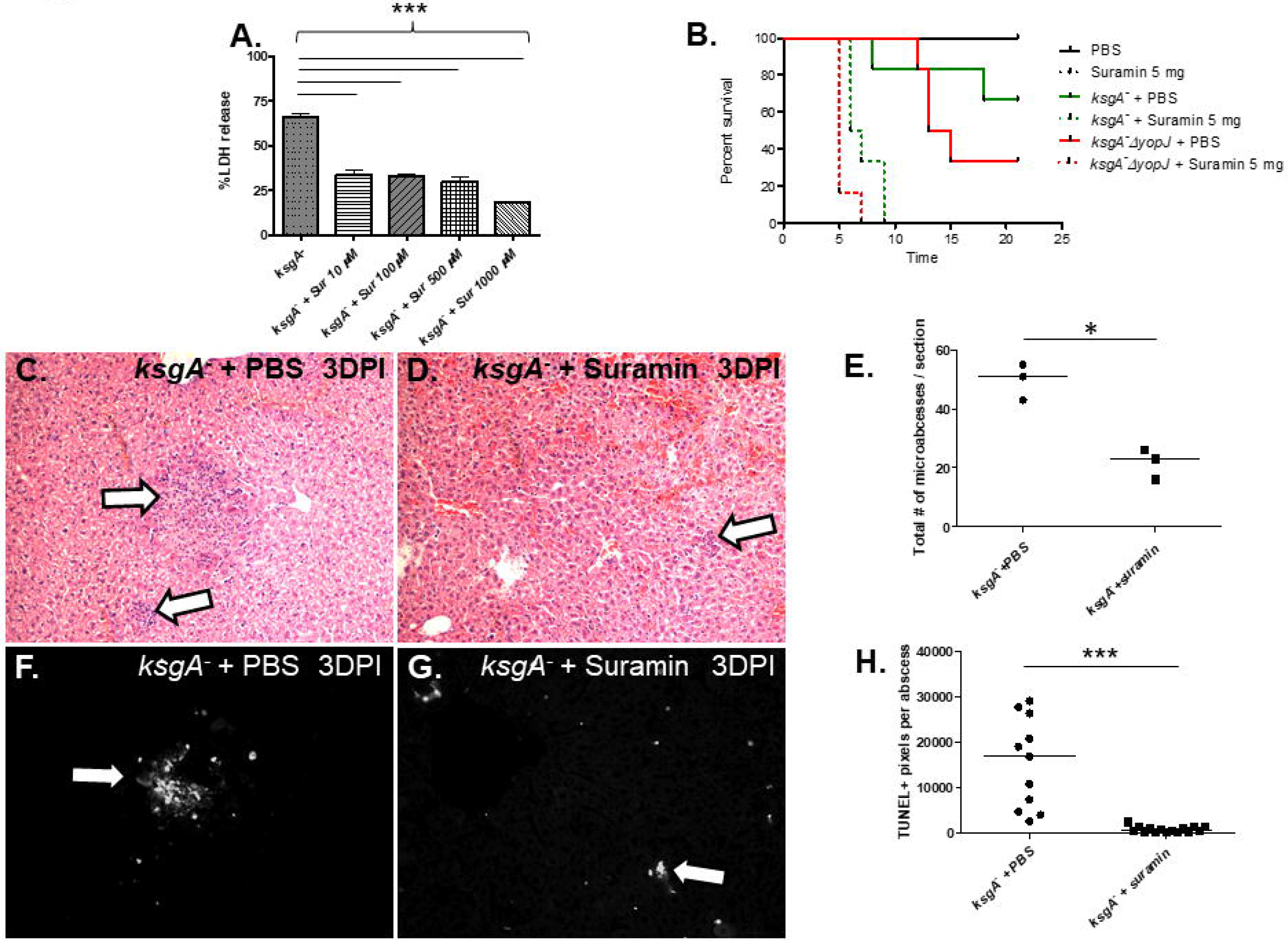
Suramin inhibits Yersina-induced macrophage cytotoxicity and enhances virulence *in vivo*. A. Bone marrow derived macrophages from C57BL/6 mice were exposed to *ksgA^-^ Y. pseudotuberculosis* at an MOI of 100:1 in the presence or absence of suramin at different concentrations. LDH release was measured to determine macrophage cytotoxicity. **B.** C57BL/6 mice (*n*=3-6/group; repeated twice) were infected intravenously (1.08∼1.38x10^3^ CFU) with various strains of *Y. pseudotuberculosis*in the presence or absence of suramin delivered intraperitoneally shortly after infection. Mice were weighed daily and monitored for signs of morbidity (*ksgA^-^* + PBS vs. *ksgA^-^* + suramin, *p*=.0042; *ksgA^-^ΔyopJ* +PBS vs. *ksgA-ΔyopJ* + suramin *p*=.0005; *ksgA-* + suramin vs. *ksgA^-^ΔyopJ* + suramin, *p*=.023). **C.** and **D.** C57BL/6 mice (*n*=3 per group; repeated twice) were infected intravenously with *Y. pseudotuberculosis ksgA-* (1x10^3^ CFU) and treated intraperitoneally with either 5 mg of suramin or with PBS. After three days, mice were sacrificed and livers were harvested and stained with H&E, large white arrows indicate representative lesions found in the sections. **E**. Quantification of the number of microabscesses from the entirety of the H&E-stained liver sections (*n*=3 per group). **F.** and **G.**, TUNEL-stained liver sections are shown, with TUNEL+ lesions highlighted with white solid arrows. **H**. Quantification of the degree of TUNEL positivity per microabscess (*n*=4-6 microabscesses per liver section; *n*=3 mice per group). The Kaplein-Meier method was used to generate survival curves and the log-rank test was used to calculate the significance. The Student’s t test was used to analyze BMDM infections. The Mann-Whitney U test was used to compare TUNEL data (\**p*<.05,\*\**p*<.01, \*\*\**p*<.005).

## Discussion

Two surprising things emerge from our study. First, positive selection of YopP/J residues has occurred directionally with *Yersinia* evolution, and has yielded an isoform with a diminished ability to induce macrophage cell death. Second, YopJ function is inversely related to bacterial pathogenesis – a hypercytotoxic YopJ variant attenuates *Yersinia* virulence in mammals, while non-functional YopP/J isoforms (or null alleles) confer hypervirulence in the context of sub-acute infection. The evolutionary forces that selected for bacteria with reduced YopJ activity or secretion are obscure, but this obscurity does not negate that positive selection of YopP/J codons has indeed occurred and these polymorphisms influence the degree of YopP/J-induced macrophage death and/or *Yersinia* virulence towards animals (**Figure 1**, see also (24, 26, 48, 49)). While we observed the YopJ phenotype in a model of sub-acute *Y. pseudotuberculosis* illness in inbred mice, we speculate that the selective niche was/is necessarily a warm-blooded wild animal. This speculation is based on the knowledge that YopJ and other Yops are not expressed at the ambient temperatures typical of fleas and soil (50, 51), the other niches known to harbor *Y. pestis (52)*, such that YopJ likely evolved to target mammalian proteins.

How is YopJ activity diminishing *Y. pseudotuberculosis* virulence? Recent evidence has shown that YopJ can block innate signaling interfering with TGF-β-activated kinase 1 (TAK1) in addition to interfering with NFκB signaling and MAPK pathways (53-56). YopJ blockade of the aforementioned signaling events and in the presence of TRIF or TNF signaling has been proposed to trigger receptor-interacting protein kinase 1 (RIPK1) and caspase-8 dependent apoptosis (55, 57, 58). Collectively, the above studies have shown that Yersinia mediated RIPK1/caspase-8 mediated apoptosis can be protective for the host. Although the precise mechanism of Yersinia induced cell death was not addressed in this study, our results similarly suggest that tailoring Yersina’s ability to induce host cell cytotoxity may have an impact on the *in vivo* pathogenesis. We further speculate that the mechanism is similar to the one we observed previously in our studies of CD8+ T cell-mediated clearance of *Y. pseudotuberculosis*-associated target cells (59). In this model, bacteria expressing ancestral or hyperfunctional YopJ attach to the host cell and inject the YopJ protein, leading to host cell death. As dead host cells are rapidly efferocytosed by neighboring activated phagocytes, the attached bacteria are also removed in the process of efferocytosis, leading to pathogen clearance (**Figure 7A**). In contrast, bacteria expressing no YopJ or catalytically dead YopJ would fail to induce host cell death, meaning the host cell would not be removed and bacteria could persist longer (**Figure 7B**).

**Fig. 7.**
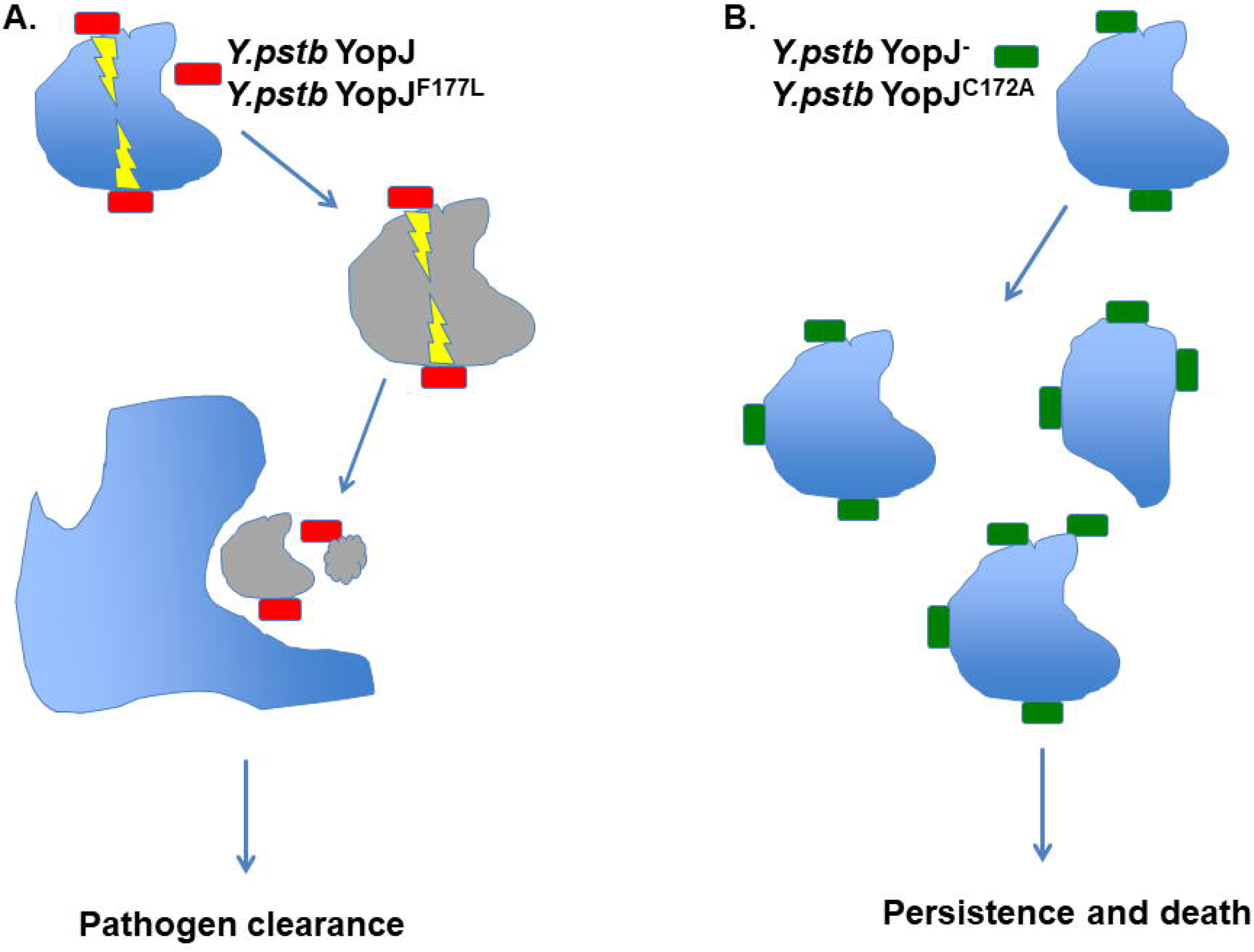
Proposed model to explain how different YopJ proteins lead to different *in vivo* outcomes following *Y. pseudotuberculosis* infection. A. *Y. pseudotuberculosis* expressing functional YopJ (harboring either the wild type/positively selected F177 or the ancestral L177) attach to the host cell and inject YopJ, leading to host cell death. Efferocytosis of dead host cells by neighboring activated phagocyte also removes those bacteria attached to dead host cells, leading to pathogen clearance **B**. *Y. pseudotuberculosis* expressing no or catalytically dead YopJ fail to induce host cell death, meaning the host cell would not be removed and bacteria could persist longer to cause illness. For either scenario, other factors including host innate and adaptive immune responses (not shown), could also influence the balance between pathogen clearance and pathogen persistence.

Given that the YopJ activity can be detrimental to *Yersinia* virulence, what then is the purpose of YopJ-induced macrophage death for *Yersinia*, when the absence of YopJ so clearly can increase *Yersinia* virulence during sub-acute illness? The continued maintenance of the *yopJ* gene in the *Y. pseudotuberculosis* and *Y. pestis* genomes, the latter of which has undergone marked decay (5, 6, 32), suggests that the existence of a niche where YopJ function remains beneficial to *Y. pestis*. One possibility is that YopJ’s beneficial or detrimental effects on bacterial virulence depend on the host tissue environment. This speculation is supported by prior observations that YopJ-null *Y. pseudotuberculosis* bacteria have mild defects in virulence via oral inoculation, but no defects via parental administration (18, 20, 24), suggesting that YopJ promotes extraintestinal dissemination of *Yersinia* or is irrelevant in systemic tissues during acute infection, a hypothesis supported by recent observations that YopJ alters intestinal permeability and promotes *Yersinia* extraintestinal dissemination (24, 60). Other findings also point to a role for YopJ in the initial local infection site (25). Additionally, the role of YopJ in *Y. pestis* has been shown to be critical for dissemination and in triggering necroptosis rather than apoptosis. This study also showed that a ΔYopJ strain of *Y. pestis* led to host survival rather than death as has been observed with ΔYopJ strains of *Y. pseudotuberculosis*(57, 61). Our findings with sub-acute systemic illness, however, demonstrate that YopJ actually hinders or impairs *Yersinia* virulence in systemic tissues. We speculate that YopJ activity serves as self-regulating mechanism for *Yersinia* pathogenesis by allowing bacterial persistence in a resistant host or tolerant tissue, possibly in a relationship approaching commensalism or at least mild parasitism. In other words, an existence comparable to that observed with other chronic illness-causing pathogens – for example, *Salmonella typhi*, which persists in gall bladders and can be shed to dissemination to new environments and hosts. Another *Yersinia* Yop can also subvert host immune recognition to promote bacterial virulence – YopK inhibits inflammasome recognition of the *Yersinia* type III secretion system and thus prevents bacterial clearance (7), although it has been proposed that YopK acts by regulating T3SS translocation, rather than by direct activity towards target proteins within the host cytoplasm (62, 63). Additionally, there is one report demonstrating that loss of YopT renders *Y. enterocolitica* hypervirulent (23). It may be that some Yops act to indirectly counter those Yops that promote virulence (i.e. YopE, YopH).

Our observations of mice exposed to non-cytotoxic *ksgA^-^* bacteria, and of mice treated with the death inhibitory chemical suramin during *Y. pseudutuberculosis* infection, strongly suggest that host cell death is beneficial for host survival and detrimental to bacterial replication and/or existence. Our experiments clearly show that suramin-treated/*ksgA^-^*-infected animals have a marked reduction in dead host cells (TUNEL^+^), suggesting that the diminished apoptosis plays at least a partial role in the increased susceptibility of suramin-treated animals to infection. Suramin has also been shown to inhibit cytotoxicity, decrease leukocyte infiltration, and dampen cytokine responses in other *in vivo* rodent models (64, 65). Therefore, it must be acknowledged that suramin has many potential uses and targets due to its complex pharmacology that may also be contributing to the results that we have observed with our Yersinia infection model (66). Our efforts to reduce host cell death levels with caspase inhibitors during *in vivo* infection were unsuccessful (data not shown), perhaps due to functional redundancy in the host cell death program components or the known consequence of increased necrosis/necroptosis following caspase inhibition (67).

YopP/J has homologues in other bacterial pathogens of animals(68), including *Vibrio parahaemolyticus* (VopP/A) (69, 70), *Aeromonas salmonicida* (AopP) [60], and *Bartonella quintana* strain Toulouse (YopP) (71). One interesting YopJ homologue is AvrA, which is found in numerous species, subspecies and serovars of the genus *Salmonella (33, 72)*. Like YopJ, AvrA is an deubiquitinase and acetyltransferase capable modifying and inhibiting MAPKK and kinases in the JNK and NFκB signaling pathways (73, 74), leading to modulation of host cell inflammatory responses in the intestine (75). Interestingly, unlike YopJ, AvrA actually prevents apoptosis in host cells (74), suggesting that sequence differences between the two proteins may explain the divergence in function. Although AvrA does possess a leucine at the position that aligns with YopJ residue 177, suggesting that AvrA should be hyperfunctional relative to WT YopJ (YopJ^F177^), prior studies have shown that AvrA affects neither cytokine expression nor plays a role in macrophage killing when expressed by either *Salmonella* or *Yersinia (76)*. However, it should be noted that the *avrA* alleles used in these studies all possess a three nucleotide deletion that results in the absence of a leucine residue at position 139 (77), also data not shown), which may explain why AvrA reportedly differs from YopJ in activity and function. However, regardless of the differences in sequence and tissue culture cell interactions, AvrA has two major features in common with YopJ – ability to attenuate virulence (74) and promote chronic infection [69] in mouse models of salmonellosis.

The ability of YopJ and AvrA to attenuate bacterial virulence is a feature typical of those YopJ homologues found in plant-associated bacteria. YopJ is a founding member of the YopJ/HopZ/AvrRxv superfamily – a family of T3SS-translocated proteins, expressed by both bacterial plant pathogens and symbionts, that are recognized by plant resistance (R) proteins during a plant defense response called the hypersensitive response (HR) (68). Induced host cell death is a critical feature of HR, and serves to limit spread of the bacterium beyond the initial infection site (78). YopJ homologues in plant-associated bacteria induce plant cell death in diverse hosts, including *Arabidopsis*, soybean, rice, tobacco and pepper plants, among others, by targeting specific plant proteins for modification (68). The requirement of a common catalytic triad suggests the plant YopJ homologues have similar enzymatic activity as YopJ. Supporting this, PopP2 from *Ralstonia solanacearum* has been shown to acetylate the *Aradopsis* RRS1-R resistance protein [71], *Xanthomonas campestris* pathovar *vesicatoria* AvrXv4 is a SUMO protease (79), and members of the HopZ family from *Pseudomonas syringae* have been shown to have protease or acetyltransferase activity (31, 80).

Another observation from our studies is reminiscent of plant R protein recognition of effectors. R proteins sense “altered self” in several ways – 1) direct modification of the R proteins by injected bacterial effector proteins, 2) indirect recognition of effector protein-driven modifications of other host proteins or 3) indirect recognition of host cell molecules released in effector translocation or bacterial infection (81). We find it intriguing that bacteria expressing YopJ^C172A^ did not phenocopy the YopJ^-^ bacteria for mouse mortality, and were <u>slightly</u> attenuated relative to the YopJ^-^ bacteria. This has been seen before with YopO/YpkA – in a study of YopO/YpkA mutant strains - *Y. pseudotuberculosis* expressing non-functional YopO was attenuated, whereas a YopO-null strain was unaffected for virulence (82). We speculate that a non-functional YopJ protein may be perceived differently by the host than either a functional YopJ or the absence of YopJ, perhaps due to host recognition of the YopJ^C172A^ protein itself or recognition of YopJ^C172A^ binding to target host proteins. This recognition would then induce a cytosolic innate immune response to the foreign (albeit inactive) YopJ protein. This speculation remains to be addressed.

Phylogenetic and phylogenomic analyses are powerful approaches to identify signatures of molecular evolution in the genes and genomes of bacterial pathogens. A hallmark paper from Ma et al. determined that the C-terminus of HopZ, the YopJ homologue found multiple *P. syringae* pathovars, has been under strong positive selection (31), providing further evidence that bacterial effectors that modulate host cell death are shaped by strong evolutionary forces. Interestingly, of the 14 pathovar HopZ sequences examined, all possessed a leucine at the position corresponding to YopJ 177 (31), suggesting that expression of a highly-functional HopZ is evolutionarily advantageous to multiple *P. syringae* pathovars. A recent phylogenomics study of 60 *Chlamydia trachomatis* strains detected signatures of positive selection in multiple T3SS effectors, and also revealed that this selection is driving *C. trachomatis* evolution towards niche-specific adaptations (colonization of particular cell types or tissues)(30). It is almost certain that global or gene-specific analyses of molecular evolution in bacterial effector proteins will reveal evidence of positive selection, even if the selective forces and niches remain obscure.

Our work extends the findings of two recent *Yersinia* phylogenomic studies of note. Gu et al. examined the evolution and divergence of multigene families from 5 *Yersinia* strains (4 *Y. pestis*, 1 *Y. pseudotuberculosis*) and found evidence of functional diversities in genes associated with pathogenicity (83). While suggestive of positive selection, the authors did not individually analyze Yops for microevolutionary signatures. In the second study, Cui et al. analyzed the genomes of 133 *Y. pestis* strains and identified 2,326 single-nucleotide polymorphisms (SNPs) that appear to have been fixed by neutral evolution rather than Darwinian selection (84). However, the authors only analyzed SNPs from the core-genome of *Y. pestis*, not the accessory genome that includes the extrachromosomal plasmid encoding the *yop* genes, meaning that *yopJ* was not included in their analysis. Others have found at least 1 chromosomally-encoded *Yersinia* protein has undergone positive selection at the codon level, as analysis of OmpF porin sequence from 73 different *Yersinia* strains found positively selected residues in several surface-exposed OmpF domains (85). As such, our findings constitute the first description of diversifying selection in the accessory genome of any *Yersinia* species member. Although our analysis were performed on only a small subset of *Yersinia* YopP/J sequences, the site identified here, codon 177, showed a significant level of positive selection, indicating the validity of our findings.

Our observations prompt the intriguing speculation that other Yops will also show evidence of molecular evolution. Given that the number of translocated *Yersinia* proteins is low (∼6-8 proteins), it seems likely that the importance of each individual effector is therefore increased, such that sequence divergence in response to any selective pressure would be greater. Microevolution within the sequences of other Yops, which generally function as pro-virulence factors, could result in Yops with increased functional abilities that confer increased virulence to individual strains or species. If true, it would present an interesting tug of war between pro-virulence Yops and anti-virulence YopP/J, the outcome of which would likely dependent on the niche.

Finally, the degree of positive selection in host proteins targeted by YopJ or other Yops is completely unknown. Recent observations of the molecular “arms race” between host and viral proteins demonstrated that the host anti-viral protein PKR (protein kinase R) has been subjected to intense episodes of positive selection in primates, in order to evade viral mimics of the PKR binding partner eIF2α (86). Regarding the latter, it is interesting to note that the activities both YopJ and YpkA/YopO converge on a kinase of eIF2α (87). Further identification of YopP/J-specific target proteins and evaluation of sequence divergence through the primate lineage will reveal the degree of YopP/J-driven evolution of host immune response proteins.

## Materials And Methods

### Bacterial Strains and Growth Conditions

Strains used in this study are indicated in Table 1, primers used for strain construction are indicated in Table 2. Lysogeny Broth (LB) and LB agar plates were made using the Lennox formulation (5g/L NaCl), and when necessary, selective antibiotics included at the following concentrations: kanamycin 30μg/mL, ampicillin 100μg/mL, irgasan 1μg/mL, chloramphenicol 20μg/mL. The *Y. pseudotuberculosis* strain YPIII pIB1 was the parent of all strains described here. The *ksgA^-^* strain has been described and used by us previously (88, 89) and contains a kanamycin-resistance-encoding transposon inserted into the *ksgA* gene. To generate the *ksgA^-^* Δ*yopJ* strain, we used a previously-described suicide vector construct, pCVD442-Δ*yopJ* (kind gift of Dr. Joan Mecsas, Tufts University), in which the upstream and downstream DNA flanking *yopJ* open-reading frame was fused together with only the first and last codon of *yopJ* remaining (90). Introduction of this unmarked *yopJ* deletion allele into the *ksgA^-^* genome was accomplished via allelic exchange, using pCVD442-encoded ampicillin and sucrose resistance as the selectable and counter-selectable markers, respectively. Briefly, the suicide vector was transferred from *Escherichia coli* strain SM10λpir to *Y. pseudotuberculosis* strain *ksgA^-^* by conjugation, and ampicillin-resistant/irgasan-resistant integrants selected. Integrants were grown overnight in the absence of plasmid selection, then dilutions of the culture placed on counter-selection agar plates containing 10% sucrose and lacking salt. Sucrose-resistant colonies were confirmed to be ampicillin-sensitive and the presence or absence of the Δ*yopJ* determined by PCR screening. To generate *yopJ* alleles carrying mutations in codons 172 and 177, a fragment of the *yopJ* gene was first cloned by PCR into pACYC184, then site-directed mutagenesis performed using long-range PCR with primers containing the mutant sequence and Dpn to remove the vector template. After confirming the sequence change, the mutated *yopJ* sequence was transferred to pCVD442, allelic exchange performed as described above, and the presence or absence of the mutant *yopJ* codons was determined by sequencing amplified genomic DNA. Bacteria were grown for cultured macrophage experiments as follows. After 2 days growth on LB agar plates from glycerol stocks, single colonies from the indicated strains were inoculated into 2 mL LB broth and incubated at 26°C for 15-18 hours with rotation. Overnight cultures were back-diluted to an estimated OD_600_ of 0.2 into low-Ca^2+^ medium (2XYT broth containing 20 mM of the Ca^2+^ chelator sodium oxalate and 20mM MgCl_2_, then grown with constant rotation for 1.5 hours at 26°C and 1.5 hours at 37°C. Cultures were then adjusted to the desired concentration in macrophage media and used for macrophage challenges as described below. Bacteria were grown for mouse experiments as follows. After 2 days growth on LB agar plates from glycerol stocks, single colonies were inoculated into 2 mL LB broth and incubated at 26°C for 15-18 hours with rotation. Overnight cultures were then adjusted to the desired concentration in PBS and used for mouse challenges as described below (after removing a sample for accurate determination of colony-forming units per mL i.e. CFU/mL).

### Codon-level selection analysis of YopJ sequences across multiple Yersinia lineages

To investigate the evolutionary history of YopJ, orthologs were found by using reciprocal blast searches across 15 sequenced genomes, including extrachromosomal elements, and 2 individual *yopJ* loci (NC_003131.gbk, NC_003132.gbk, NC_003134.gbk, NC_003143.gbk, NC_004088.gbk, NC_004838.gbk, NC_005810.gbk, NC_005813.gbk, NC_005814.gbk, NC_005815.gbk, NC_005816.gbk, NC_006153.gbk, NC_006154.gbk, NC_006155.gbk, NC_008118.gbk, NC_008119.gbk, NC_008120.gbk, NC_008121.gbk, NC_008122.gbk, NC_008149.gbk, NC_008150.gbk, NC_008791.gbk, NC_008800.gbk, NC_009377.gbk, NC_009378.gbk, NC_009381.gbk, NC_009704.gbk, NC_009705.gbk, NC_009708.gbk, NC_010157.gbk, NC_010158.gbk, NC_010159.gbk, NC_010465.gbk, NC_010634.gbk, NC_010635.gbk, NC_014017.gbk, NC_014022.gbk, NC_014027.gbk, NC_014029.gbk, NC_015224.gbk, NC_015475.gbk, NC_017153.gbk, NC_017154.gbk, NC_017155.gbk, NC_017156.gbk, NC_017157.gbk, NC_017158.gbk, NC_017159.gbk, NC_017160.gbk, NC_017168.gbk, NC_017169.gbk, NC_017170.gbk, NC_017263.gbk, NC_017264.gbk, NC_017265.gbk, NC_017266.gbk, NC_017564.gbk, NC_017565.gbk).

Protein sequences were aligned using Clustalw and nucleotide sequences for each YopJ ortholog were aligned based on their amino acid translation (using in house scripts). These nucleotide alignments were used as input to RAxML[83], where maximum likelihood phylogenies were generated (GTRGAMMA model of substitution). A considerable amount of change in the YopJ homologs has occurred during the evolution of these pathogens; some of these changes alter protein encoding (dN, non-synonymous nucleotide substitutions) while some do not (dS, synonymous substitutions). Evidence of adaptative selection was found across the aligned ortholog set by calculating the ratio of dN/dS in pairwise comparisons of orthologs. In order to identify those YopJ residues under selection and the direction and strength of selection at the codon level, the frequency of nonsynonymous and synonymous changes and their ratio (dN/dS or ω) were calculated using Phylogenetic Analysis by Maximum Likelihood (PAML v4.7) software using the Goldman and Yang amino acid substitution model (codeML) and these same nucleotide alignments [84]. To increase the sensitivity of our analysis we also calculated variable dN/dS (*ω*) over all sites (models M7 vs. M8) and used the likelihood ratio test to identify statistically significant evidence for diversifying selection (χ2 = 14; df = 2; p < 0.01). In house Perl scripts were used to parse the codeML output files to identify residues predicted to have undergone diversifying selection in YopJ (using maximum likelihood ratio tests before analyzing the results).

### Derivation of Bone Marrow Macrophages

Bone marrow-derived macrophages (BMDM) were generated by culturing bone marrow cells freshly-harvested from the femurs and tibias of C57BL/6 mice in supplemented DMEM as previously described (59). Macrophages were harvested on day 6 of the derivation protocol and seeded at a density of 5 x 10^4^ cells/100μL/well of a 96 well plate in supplemented DMEM lacking antibiotics and allowed to adhere to the plastic overnight. Occasionally, at day 6, macrophages were cryofrozen in 5% DMSO/95% fetal-bovine serum (FBS) at a concentration of 1x10^7^ cells/mL and subsequently thawed for seeding.

### Macrophage Infections, Treatments and Cytotoxicity Assay

*Y. pseudotuberculosis,* grown as described above, was added to adherent macrophages at a multiplicity of infection (MOI) of 100:1 in DMEM without phenol red + 1% FBS for 45 minutes to 1 hour. Gentamicin was added to a final concentration of 100 μg/mL and the cells were allowed to incubate an additional 2 hours. For suramin experiments, BMDM were first pretreated with various concentrations of suramin (Sigma, St. Louis, MO) for 1 hour, then exposed to bacteria at an MOI of 100:1 in the presence or absence of suramin at the indicated concentrations. Macrophage death was evaluated by detection of the cytoplasmic enzyme lactate dehyrogenase (LDH) in the culture supernatants similarly as previously described(59). Briefly, macrophages were spun down at 250xg for 2 minutes and 50μL supernatant carefully removed from each well, transferred to a new 96-well plate, and immediately assayed for the presence of LDH using the Cytotox 96 Assay kit (Promega, Madison, WI) according to the manufacturer’s recommendations. Note that controls cells were treated with 1% Triton-X 100 to liberate cytoplasmic contents approximately 15 minutes prior to supernatant harvest. 50μL of reconstituted substrate solution (containing 2-(4-Iodophenyl)-3-(4-nitrophenyl)-5-phenyl-2H-tetrazolium chloride, or INT) was added to the harvested supernatants, the reaction incubated at room temperature in the dark for 30 minutes, then the enzymatic conversion of INT into a red formazan product terminated by addition of 50μL stop solution (1M acetic acid) and excess bubbles in each well punctured. Absorbance was measured at 490 nm in a BioTek Synergy H4 plate reader (BioTek, Winooski, VT). % Cytotoxicity was calculated as follows: 100× [(experimental release – effector T cell spontaneous release – target cell spontaneous release)/(target cell maximum release – target cell spontaneous release)].

### Mouse Infections and Treatments

6-8 week old female C57BL/6 mice were purchased from the National Cancer Institute colony at Charles River. Mice were allowed to rest for 7 days after arrival, then infected intravenously via tail vein injection with a 200 μL inoculum containing between 3.5 x 10^2^ and 2 x 10^3^ colony-forming units (CFU) in PBS of the indicated *Y. pseudotuberculosis* strain. Mice were then visually inspected for signs of illness including lethargy and piloerection and weighed daily to monitor the disease progression. For suramin experiments, mice were infected as before, then intraperitoneally injected once with 5 mg suramin/400 μL PBS or 400 μL PBS alone 30 minutes to 1 hour post-infection. For clodronate experiments, clodronate-containing liposomes were prepared as described below, then 200 μl of liposomes were injected intravenously 6 hours post infection. All animal experiments were performed in accordance with the NIH Guide for the Care and Use of Laboratory Animals and were approved by the UTHSCSA Institutional Animal Care and Use Committee.

### Preparation and Characterization of Clodronate Liposomes

Liposomes containing dichloromethylene bisphosphonate (clodronate) or control empty liposomes containing phosphate buffered saline (PBS) were prepared from reported protocols with slight modifications(45, 91). In two 500 mL round bottom flasks, 96 mg cholesterol (Calbiochem, San Diego, CA) was added and dissolved with 10 mL chloroform (HPLC grade, Fisher Scientific, Fair Lawn, NJ). Then 4.14 mL of egg phosphatidylcholine in chloroform (25 mg/ mL; Avanti Polar Lipids, Alabaster, AL) was added. Lipid films were formed following removal of chloroform by rotary evaporation. The lipid films were placed under vacuum for 1 h to remove residual chloroform. The lipid films were rehydrated with 12 mL of 0.7 M clodronate (Sigma Aldrich, St Louis, MO) prepared in degassed sterile water for injection (Hospira, Lake Forest, IL) and pH adjusted to 7.0, or degassed Dulbecco’s PBS, pH 7.4 (Invitrogen, Carlsbad, CA). The flasks were swirled until the lipids were in suspension. The lipid suspensions were kept at room temperature for 2 h with occasional swirling, sonicated in a water bath for 4 minutes (Branson, Danbury, CT) and then kept at room temperature for 2 h. The liposome samples were then extruded through polycarbonate filters (2 µ, 2 times; 1 µ, 4 times) using an extruder (Lipex, Northern Lipids, Vancouver, Canada) and collected in a new tube. After flushing the liposome samples with argon, the samples were stored overnight at 4° C. The next day the liposomes were transferred to microcentrifuge tubes and centrifuged in an Allegra 21R centrifuge (Beckman Coulter, Brea, CA) at 15,300 rpm for 30 minutes at 10°C. The supernatants were removed and the pellets resuspended with degassed PBS. The samples were washed 4 times. After the last wash, the liposome pellets were resuspended in degassed PBS to a final volume of 4.8 mL, flushed with argon and stored at 4°C until needed. Liposomes were used for the study within 2 weeks of preparation. Liposome diameter was measured at 488 nm with a DynaPro dynamic light scattering system (Wyatt Technology, Santa Barbara, CA). The diameters for clodronate liposomes and control PBS liposomes were 968 ± 28 nm and 790 ± 41 nm, respectively. Phospholipid content was measured using phosphatidylcholine colorimetric assay kit (Cayman Chemical, Ann Arbor, MI) and determined to be 14.5 mg/mL for the clodronate liposomes and 15.4 mg/mL for the PBS liposomes. The amount of liposome-encapsulated clodronate was determined to be 12.7 mg/mL (45). In pilot experiments, clodronate liposomes were confirmed to induce macrophage death by TUNEL staining in spleens of non-infected mice (data not shown).

### Organ Bacterial Burden Assay

At the noted times post-infection, mice were sacrificed by cervical dislocation, spleens and livers aseptically removed, transferred to sterile PBS from mice at various time points after infection and weighed. Tissues were mechanically disrupted using a tissue homogenizer (Omni, Marietta GA), serially diluted in sterile PBS and 100 μL of the dilutions plated on nutrient LB agar plates. After 48 hours at 26°C, bacterial colonies were enumerated and CFU normalized to weight of tissue in grams. Note that as each animal may have had an organ weight different from the other animals, the CFU values at or below the limit of detection (10 CFU per mL of organ homogenate) were occasionally different between individual animals. The limit of detection in the spleen was (90 CFU/g/spleen or 1.95 Log CFU) and in the liver was (22 CFU/g/Liver or 1.33 Log CFU).

### Cytokine Analysis

Cytokine analysis was performed on serum and tissue homogenates from day 3 and day 9 post-inoculation. Briefly, blood was collected via cardiac puncture and allowed to clot for 30 minutes. Serum was collected after centrifugation at 7000 rpm for 7 minutes and stored at -80°C. Cytokines were assayed using the Mouse Cytokine 20-plex panel strictly adhering the manufacturer’s protocol (Life Technologies, Carlsbad CA). Cytokine levels were detected using a Bio-Plex 200 system and analyzed using Bio-Plex Manager 6.0 software (Bio-Rad, Hercules CA).

### Terminal Deoxynucleotidyl Transferase dUTP Nick End Labeling (TUNEL) Assay

Spleens were harvested at 3 days post-infection and were snap-frozen in the cyrostat sectioning medium Tissue-Tek® O.C.T.™ (Sakura, Torrance CA). Livers were removed and placed in 10% buffered formalin for at least three days and then sent to the UTHSCSA Histopathology Laboratory to be embedded in paraffin and cut into 4 micron sections. Some slides with liver sections were stained with hematoxylin and eosin and others were processed for TUNEL staining with the ApopTag kit as outlined below. OCT-frozen spleen sections were cut with a cryostat (Thermo Scientific Shandon, Waltham MA), placed on slides and allowed to air-dry overnight. The slide-affixed tissue sections were then fixed in acetone for 20 seconds at room temperature, allowed to dry, then wrapped in foil and stored at -80°C. Subsequently, sections were stained for nicked DNA, to label TUNEL^+^ cells, using the ApopTag kit with included buffers (EMD Millipore, Billerica MA). Briefly, slides were removed from -80°C and allowed to warm up to room temperature. Slides were then fixed in 1% paraformaldehyde for 10 minutes at room temperature and washed in PBS twice. Tissue sections were not allowed to dry after this point. Tissue sections were then circled with a hydrophobic PapPen (Kiyota International, Elk Grove Village, IL) and equilibration buffer was incubated on the section for at least 10 seconds. TdT enzyme solution (10% enzyme + 20% PBS + 70% reaction buffer) was applied for 60 minutes in a humidity chamber at 37°C. Stop solution was then applied for 10 minutes at room temperature followed by three washes with PBS. A solution containing anti-digoxigenin fluorescein antibody (53% blocking solution + 47% anti-digoxigenin fluorescein conjugate) was applied and allowed to incubate for 30 minutes in a dark humidity chamber. Slides were then protected from light as much as possible from this point forward. Slides were then washed three times in PBS and mounted with ProLong Gold Antifade reagent containing DAPI (Life Technologies, Carlsbad CA). Sections from a given experiment were stained on the same day.

### Microscopy and Image Capture, Analysis

Stained tissue sections were visualized using either an EC plan neofluar 10x or Plan-Apochromat 20x/.8 objective on a Zeiss AxioImager Z1 epifluorescent microscope (Carl Zeiss, Thornwood, NY). Images were captured using a Zeiss AxioXam MRm Rev3 and/or MRc cameras and analyzed using Zeiss AxioVision release 4.5 software. Sections from a given experiment were imaged on the same day with the same exposure times for image acquisition to ensure consistency between samples. ImageJ (92) was used to measure the pixel intensity in each captured image.

### Data and Statistical Analysis

Prism 5 (GraphPad Software, La Jolla CA) was used for graphing and statistical analysis. Survival curves were estimated using the Kaplan Meier method and significance calculated using the log-rank test. The nonparametric Mann-Whitney U test and unpaired Student’s t test were used to determine statistical differences between groups of data from animal and tissue culture experiments, respectively. Image quantification data was evaluated for significance using the Mann-Whitney U test.

## Acknowledgements

We thank Ian Morris and Michael Berton for use and instruction of the Zeiss AxioImager Z1 microscope. We thank Joan Mecsas, Peter Dube and Angel Cantwell for helpful discussions and critical analysis of the manuscript. This work was supported by National Institutes of Health, Grant AI085116 from the NIAID to M.A.B., and Grant AI110684 from the National Institute of Allergy and Infectious Diseases and HHMI support of R.R.I.

