## Supplemental Tables and Figures for "Heightened virulence of *Yersinia* is associated with decreased function of the YopJ protein"

#### SUPPLEMENTAL FIGURE LEGENDS

**S1. Repair of *yopJ in the ksgA⁻ΔyopJ* strain restores the original phenotype. A.** RAW macrophages were infected with either *ksgA*$^{-}$, *ksgA*$^{-}$*ΔyopJ*, or *ksgA*$^{-}$*yopJ*$^{repaired}$ *Y. pseudotuberculosis* strains at an MOI of 100:1. LDH release was measured to determine the cytotoxicity of these strains in macrophages. (**B**. and **C.**) C57BL/6 mice were intravenously challenged with *ksgA*$^{-}$, *ksgA*$^{-}$*ΔyopJ* or *ksgA*$^{-}$*ΔyopJ*$^{repaired}$ *Y. pseudotuberculosis*. Morbidity and mortality was followed after infection and data shown represents the percent survival and weight loss of each group (*n*=5 mice per group).

**S2. A cytokine storm does not characterize illness caused by *ksgA-ΔyopJ* bacteria.** C57BL/6 mice were inoculated intravenously (1~1.26x10$^{3}$ CFU) with *ksgA*$^{-}$, *ksgA*$^{-}$*ΔyopJ*, or *ksgA*$^{-}$ *yopJ*$^{C172A}$ *ksgA*$^{-}$ *yopJ*$^{F177L}$ bacteria. Serum was collected from each mouse at day 9 post inoculation and was assayed for cytokines using a cytokine multiplex assay.

**S3. Systemic clodronate liposome induced macrophage apoptosis in vivo does not protect mice from *Y. pseudotuberculosis* infection.** 6-8 week old female C57BL/6 mice were infected intravenously (1x10$^{3}$ CFU) with *ksgA*$^{-}$*ΔyopJ Y. pseudotuberculosis* and left untreated or treated with either PBS containing liposomes or clodronate containing liposomes at 6 hours post infection. Morbidity and mortality was followed after infection and data shown represents the percent survival of each group (*n*=6 mice per group). The Kaplein-Meier method was used to generate survival curves and the log-rank test was used to calculate the significance (*ksgA*$^{-}$*ΔyopJ* vs. *ksgA*$^{-}$*ΔyopJ* + PBS, n.s.;

*ksgA*$^{-}$*ΔyopJ* vs. *ksgA*$^{-}$*ΔyopJ* + clodronate, *p*=.064; *ksgA*$^{-}$*ΔyopJ* + PBS vs. *ksgA*$^{-}$*ΔyopJ* + clodronate, *p*=.065).

**Table 1.** ***Y. pseudotuberculosis*** **(YPIII pIB1) strains used in this study**

| Strain | Strain name | Reference |
|---|---|---|
| *ksgA*$^{-}$ | 500 | [80] |
| *ksgA*$^{-}$*ΔyopJ* | MB153 | This study |
| *ksgA*$^{-}$*yopJ*$^{C172A}$ | MB219 | This study |
| *ksgA*$^{-}$*yopJ*$^{F177L}$ | MB254 | This study |

**Supplementary Table 1. Bayes Empirical Bayes results for positively selected sites in YopJ (*: P>95%; **: P>99%)**

| position | Residue | Prob (ω > 1) | Mean ω | Standard deviation |
|---|---|---|---|---|
| 10 | I | 0.544 | 5.537 | 4.754 |
| 11 | S | 0.998** | 9.843 | 1.001 |
| 20 | S | 0.938 | 9.291 | 2.394 |
| 33 | T | 0.790 | 7.904 | 3.852 |
| 40 | S | 0.581 | 5.891 | 4.715 |
| 52 | M | 0.999** | 9.851 | 0.965 |
| 54 | V | 0.791 | 7.920 | 3.841 |
| 80 | L | 0.989 | 9.765 | 1.311 |
| 130 | A | 0.835 | 8.324 | 3.532 |
| 139 | M | 0.740 | 7.435 | 4.128 |
| 143 | R | 0.909 | 9.020 | 2.795 |
| 177 | F | 0.971 | 9.592 | 1.798 |
| 205 | G | 0.838 | 8.356 | 3.504 |
| 206 | E | 1.000** | 9.861 | 0.916 |
| 212 | D | 0.806 | 8.059 | 3.742 |

**Supplementary Table 2. Naïve Empirical Bayes results for positively selected sites in YopJ (*: P>95%; **: P>99%)**

| position | Residue | Prob (ω > 1) | Mean ω |
|---|---|---|---|
| **10** | I | 0.852 | 3.09 |
| **11** | S | 0.998** | 3.604 |
| **20** | S | 0.727 | 2.649 |

| | | | |
|---|---|---|---|
| **33** | T | 0.749 | 2.727 |
| **40** | S | 0.894 | 3.238 |
| **52** | M | 0.999** | 3.611 |
| **54** | V | 0.759 | 2.76 |
| **55** | E | 0.742 | 2.701 |
| **62** | I | 0.833 | 3.022 |
| **75** | L | 0.791 | 2.874 |
| **80** | L | 0.888 | 3.216 |
| **95** | R | 0.765 | 2.783 |
| **106** | G | 0.802 | 2.913 |
| **130** | A | 0.79 | 2.869 |
| **139** | M | 0.686 | 2.503 |
| **143** | R | 0.83 | 3.011 |
| **144** | T | 0.776 | 2.822 |
| **177** | F | 0.995** | 3.596 |
| **185** | I | 0.859 | 3.116 |
| **189** | S | 0.846 | 3.067 |
| **205** | G | 0.79 | 2.871 |
| **212** | D | 0.674 | 2.459 |
| **242** | G | 0.825 | 2.994 |
| **243** | V | 0.962* | 3.479 |
| **244** | G | 0.839 | 3.043 |
| **245** | T | 0.998** | 3.607 |

| | | | |
|---|---|---|---|
| **246** | V | 0.962* | 3.479 |
| **247** | V | 0.931 | 3.368 |
| **248** | N | 0.979* | 3.538 |
| **251** | N | 0.990** | 3.58 |
| **252** | E | 0.699 | 2.547 |
| **253** | T | 0.861 | 3.122 |
| **254** | I | 0.999** | 3.611 |
| **255** | V | 0.999** | 3.608 |
| **256** | N | 0.912 | 3.302 |
| **257** | R | 1.000** | 3.612 |
| **258** | F | 0.793 | 2.88 |
| **259** | D | 0.998** | 3.607 |
| **260** | N | 0.979* | 3.538 |
| **261** | N | 0.912 | 3.302 |
| **262** | K | 0.66 | 2.409 |
| **263** | S | 0.885 | 3.206 |
| **264** | I | 0.992** | 3.584 |
| **265** | V | 1.000** | 3.613 |

#### Supplementary Table S3. Primers used in this study

| Primer | Sequence | Purpose |
|---|---|---|
| F yopJ | 5'CAACAAGTTTCTCTACCGGAGAAT3’ | screen for Δ*yopJ* allele |
| R yopJ | 5'CTCATACCACCCGTACTCTAGCA3’ | |
| yopJ F3 | 5’GATCGATATCCAAGTGCCCCCTAAGCCTTGAGTT3’ | PCR clone *yopJ* orf into pACYC184, screen for *yopJ* point-mutant alleles |
| yopJ R2 | 5’GATCGTCGACCCCATACTGGAGCAAGATTTCC3’ | |
| yopJ* F (sense) | 5’GAAATGGATATTCAGCGAAGCTCATCTGAAGCTGGTATTTTTAGTTTTGCAC3’ | mutate $yopJ^{WT}$ to $yopJ^{C172A}$ |
| yopJ* R (antisense) | 5’GTGCAAAACTTAAAATACCAGCTTCAGATGAGCTTAGCTGAATATCCATTTC3’ | |
| yopJ F4 | 5’GATCGCATGCCAAGTCCCCCCTAAGCCTTGAGTT3’ | PCR clone $yopJ^{C172A}$ into pCVD442 |
| yopJ R4 | 5’GATCGCATGCCCCATACTGGAGCAAGATTTCC3’ | |
| F177A F (sense) | 5’GAAGCTCATCTGAATGTGGTATTTTTAGTTTGGCACTGGCAAAAAAAC3’ | mutate $yopJ^{C172A}$ to $yopJ^{F177L}$ |
| F177A R (antisense) | 5’GTTTTTTTGCCAGTGCCAAACTAAAAATACCACATTCAGATGAGCTTC3’ | |

[1]underlined sequence indicates restriction enzyme site used for cloning

### Supplementary Figure 1

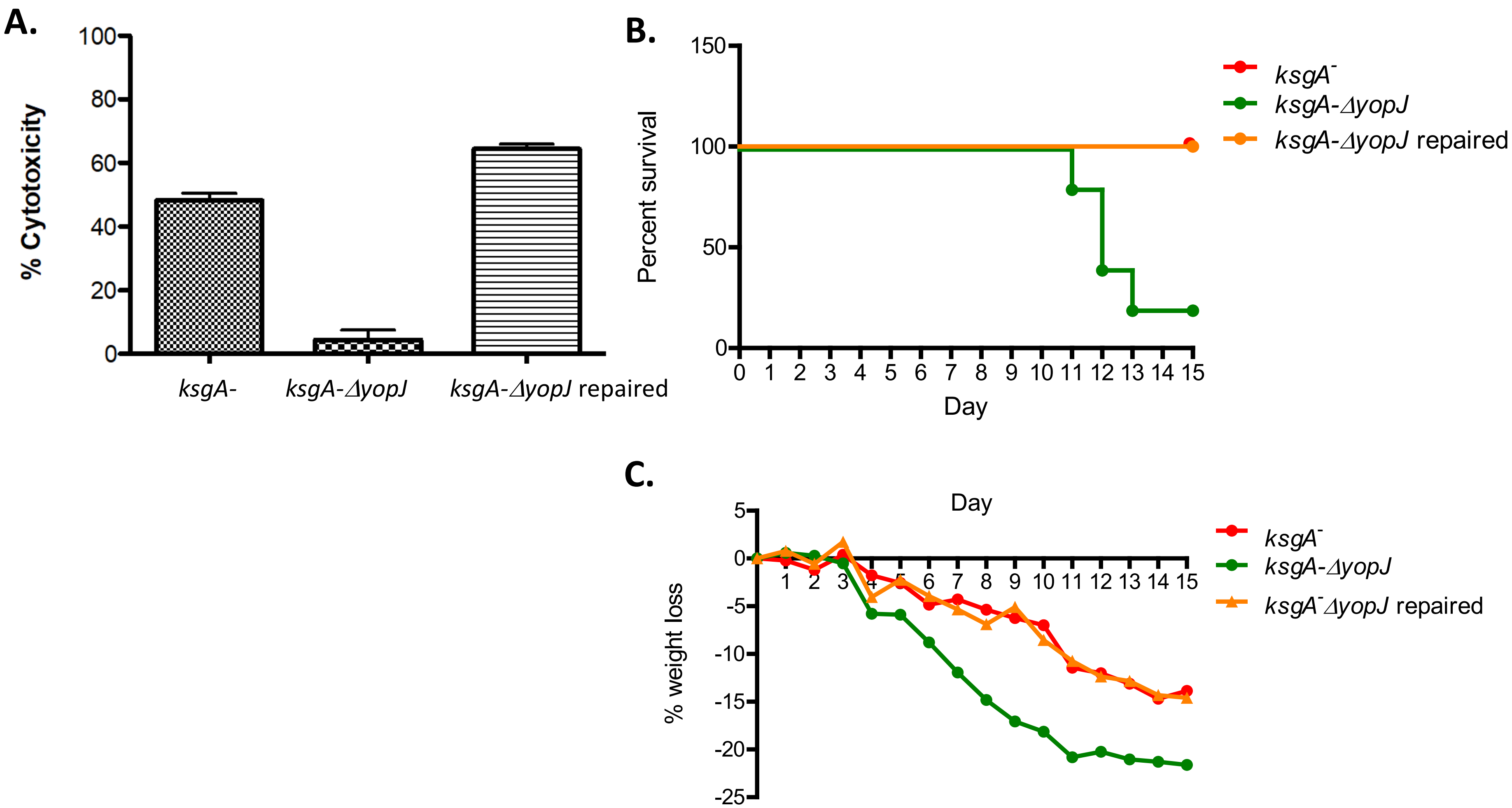

### Supplementary Figure 2

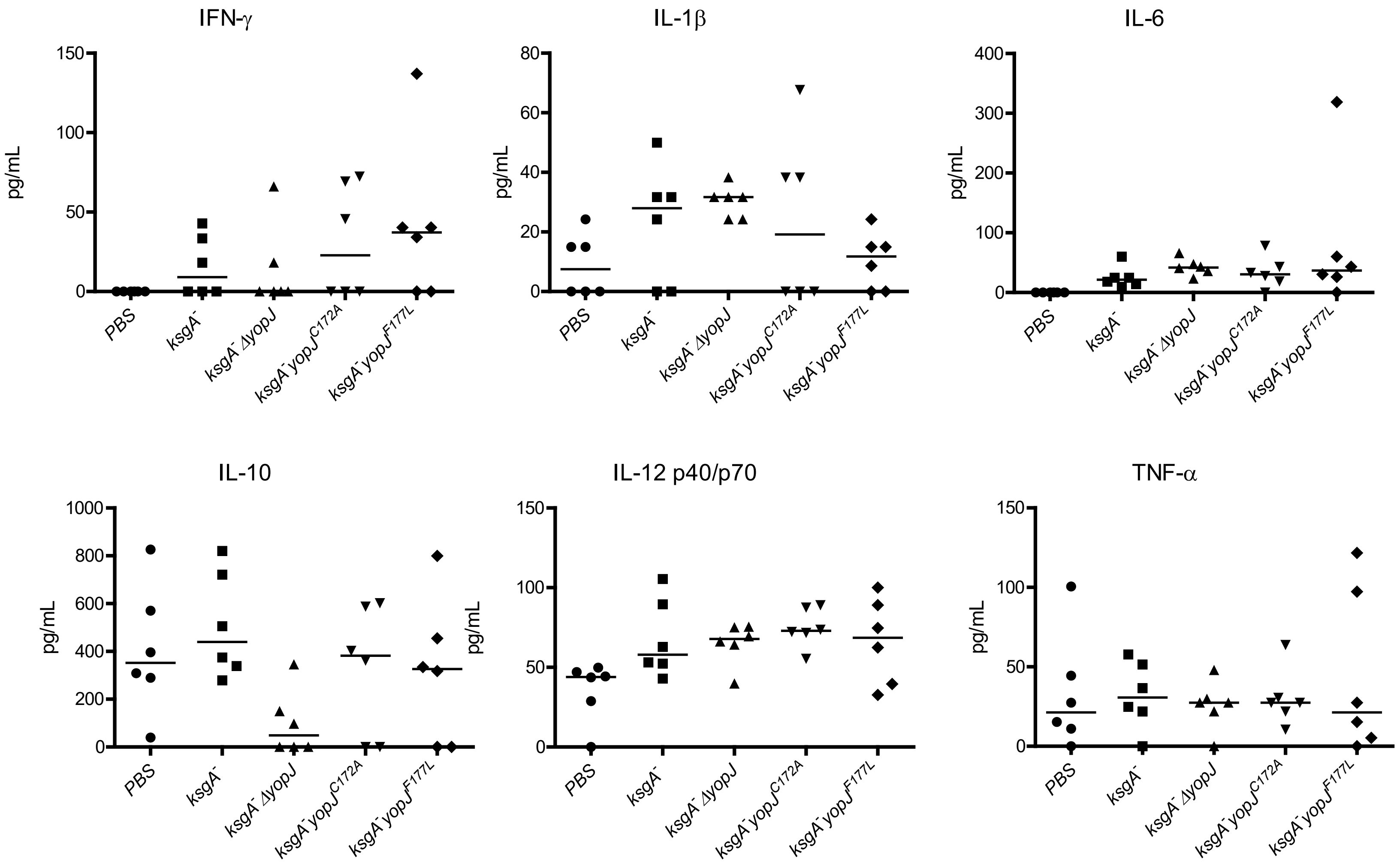

### Supplementary Figure 3

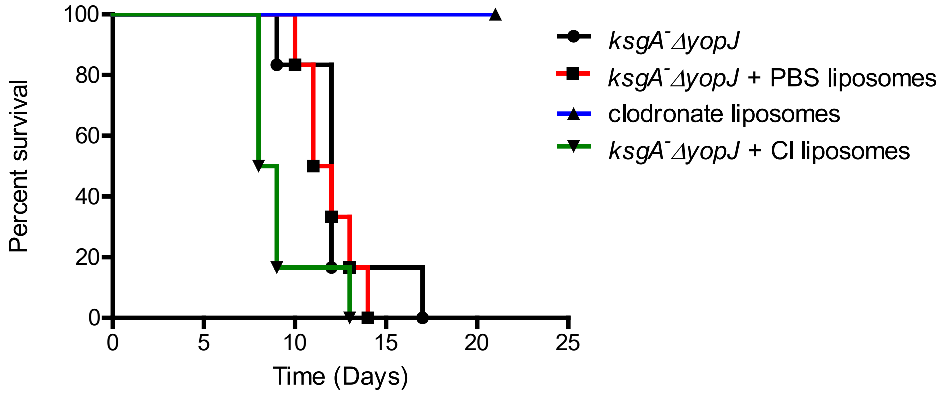
